# PuBliCiTy: Python Bioimage Computing Toolkit

**DOI:** 10.1101/2021.03.01.432926

**Authors:** Marcelo Cicconet

## Abstract

The Python Bioimage Computing Toolkit (PuBliCiTy) is an evolving set of functions, scripts, and classes, written primarily in Python, to facilitate the analysis of biological images, of two or more dimensions, from electron or light microscopes. While the early development was guided by the goal of replacing an existing internal code-base with Python code, the effort later came to include novel tools, specially in the areas of machine learning infrastructure and model development. The toolkit is built on top of the so-called *python data science stack*, which includes *numpy, scipy, scikit-image, scikit-learn*, and *pandas*. It also contains some deep learning models, written in TensorFlow and PyTorch, and a web-app for image annotation, which uses Flask as the web framework. The main features of the toolkit are: (1) simplifying the interface of some routinely used functions from underlying libraries; (2) providing helpful tools for the analysis of large images; (3) providing a web interface for image annotation, which can be used remotely and on tablets with pencils; (4) providing machine learning model implementations that are easy to read, train, and deploy – written in a way that minimizes complexity for users without a computer science or software development background. The source code is released under an MIT-like license at github.com/hms-idac/PuBliCiTy. Details, tutorials, and up-to-date documentation can be found at the project’s page as well.

**Project page:** github.com/hms-idac/PuBliCiTy

## 1 Overview

PuBliCiTy’s code is divided in 5 main areas:

- general purpose functions
- 2D and 3D Random Forest-based segmentation tools
- 2D and 3D Deep Learning-based segmentation tools
- classes to help processing large 2D and 3D images
- web-app for 2D and 3D image visualization and annotation

### 1.1 General Purpose Functions

At the time of writing, the file `gpfunctions.py` contains about 100 functions with diverse purposes, such as I/O (e.g. *imread, imwrite, writeTable*), format conversion (e.g. *rgb2gray, im2double*), image processing (e.g. *imadjustgamma, imadjustcontrast, normalize*), geometric processing (e.g. *imrotate, imtranslate*), morphological operations (e.g. *imerode, imdilate*), filtering (e.g. *medfilt, imgaussfilt, imlogfilt*), feature computation (e.g. *localstats, imderivatives*), convolutions (e.g. *circleKernel, morletKernel, conv2*), visualization (e.g. *imshow, changeViewPlane*), segmentation/detection (e.g. *findSpots2D, thrsegment, imbinarize*), among others.

Most of these functions are used routinely in image analysis tasks, while some provide the back-end to others tools - e.g. *bwInterpSingleObjectMasks*, which is used behind the scenes when annotating 3D objects via the web-app.

### 1.2 Random Forest-Based Segmentation

Certain image segmentation tasks are not so complex as to require the latest machine learning models. Nuclei segmentation is often one such case, where, if the input signal has decent signal-to-noise ratio, a “classic” machine learning model suffices.

The files `pixelclassifier.py` and `voxelclassifier.py` provide tools for 2D and 3D semantic segmentation via Random Forests. The models compute “edge” and “blob” features at scales specified by the user, and the training process outputs measurements of feature importance, so that the user can inspect which ones are more useful.

### 1.3 Deep Learning-Based Segmentation

PuBliCiTy ships with custom implementations of the U-Net [5] architecture for semantic segmentation, for 2D and 3D images (files `unet2D.py` and `unet3D.py`, respectively). These deep learning models are built to solve the same problem as the random forest models above, however they in general perform better, at the expense of (1) requiring more training data and (2) being harder to train. One design decision in developing these was to make the code as simple and readable as possible. Thus `unet2D.py` and `unet3D.py` are single, linear, highly commented scripts.

The toolkit also includes a PyTorch Mask-RCNN [1] model for instance segmentation, customized for cell (or, more generally, single object) segmentation. While the U-Net models were developed from scratch, the Mask-RCNN is based on available PyTorch architecture, pre-trained on the COCO instance-segmentation dataset [4]. From a user’s perspective, the main difference between the Mask-RCNN and U-Net models is that the output of the U-Net are pixel classes (which have to be post-processed to obtain object masks), while the Mask-RCNN predicts object label masks directly. The model is implemented in `cellMaskRCNN.py`.

### 1.4 Partition of Image Classes

Certain image analysis algorithms have restrictions on image size. For example: a deep learning architecture implemented with convolutions where the output size equals the input size cannot be deployed on images of size different from those in the training set. Or perhaps an image is too big to fit in GPU memory. Or maybe the prediction model can only detect up to 100 objects, and the image could have more.

In such scenarios, one needs to partition the input image in small portions, possibly with overlap, process these portions, then assemble the output. PuBliCiTy has three “partition of image” classes for that purpose. `PartitionOfImage.py` splits images with overlapping tiles, and performs the assembly averaging values where tiles intersect. `PartitionOfImageVC.py` splits images similarly, but only assembles the inner, non-overlapping part of the tiles. Finally, `PartitionOfImageOM.py` splits images into overlapping tiles and assembles object masks and bounding boxes, using object intersection to resolve redundancy for objects that fall into overlapping areas.

`unet2D.py, unet3D.py`, and `cellMaskRCNN.py` use these classes in “deployment” mode.

### 1.5 PuBliCiTy Web-App

An essential part of developing machine learning models is gathering annotations. In image analysis tasks this usually implies drawing areas in images which correspond to various classes of pixels (say background versus cell of interest), or drawing contours around objects. Some of these tasks can be done using existing software (e.g. Fiji), but for others we did not find easily available solutions. For example, 3D segmentation models require solid 3D annotations, not simply a sparse set of annotated planes.

Therefore we implemented our own annotation (and visualization) tools as a web-app. This architectural choice has two main advantages: first, a web interface allows annotating images remotely (i.e. in a workstation different from the one where the data is), which is useful for large image files; second, annotations can be done on tablets with pencil tools, which greatly improves efficiency.

The web-app’s main file is `publicityWebApp.py`.

## 2 Sample Applications

While PuBliCiTy includes implementations of algorithms available elsewhere, it also contains novel tools that extend the capability of those methods, and interfaces that simplify image analysis tasks. In this section we describe two such cases: (1) how `PartitionOfImageOM.py` can be used to allow a Mask-RCNN model to work on images of size much larger than it was originally designed for; (2) how `publicityWebApp.py` can be deployed to annotate images on an iPad with an Apple Pencil, allowing for faster annotations w.r.t. using a mouse on a regular computer.

### 2.1 Partition of Image

Mask-RCNN models for instance segmentation are typically optimized for images of a certain size range, due primarily to the sizes of images in available training datasets. The implementation available with PyTorch, for example, resizes by default images that have sides outside the range [800,1333]. As a consequence, even though the original model can - assuming enough compute capacity - perform prediction outside that range, results degrade.

To show this, we fine-tuned a Mask-RCNN model (with ResNet-50[2] FPN[3] backbone, pre-trained on the COCO dataset[4]) on 100 images of size 1000^2^, containing 100 cells each, then deployed the model on similar square images of side 1000, 2000, and 4000, with and without using Partition of Image.

Using the original model, while results for the image of side 1000 are near perfect, prediction quality deteriorates substantially for the image of side 2000 (see Figure 1), and accuracy is near 0 for the image of side 4000. Using Partition of Image, however, we are able to split the larger images in patches of size near optimal, and accuracy does not degrade substantially. Table 1 shows more detailed results.

**Figure 1:**
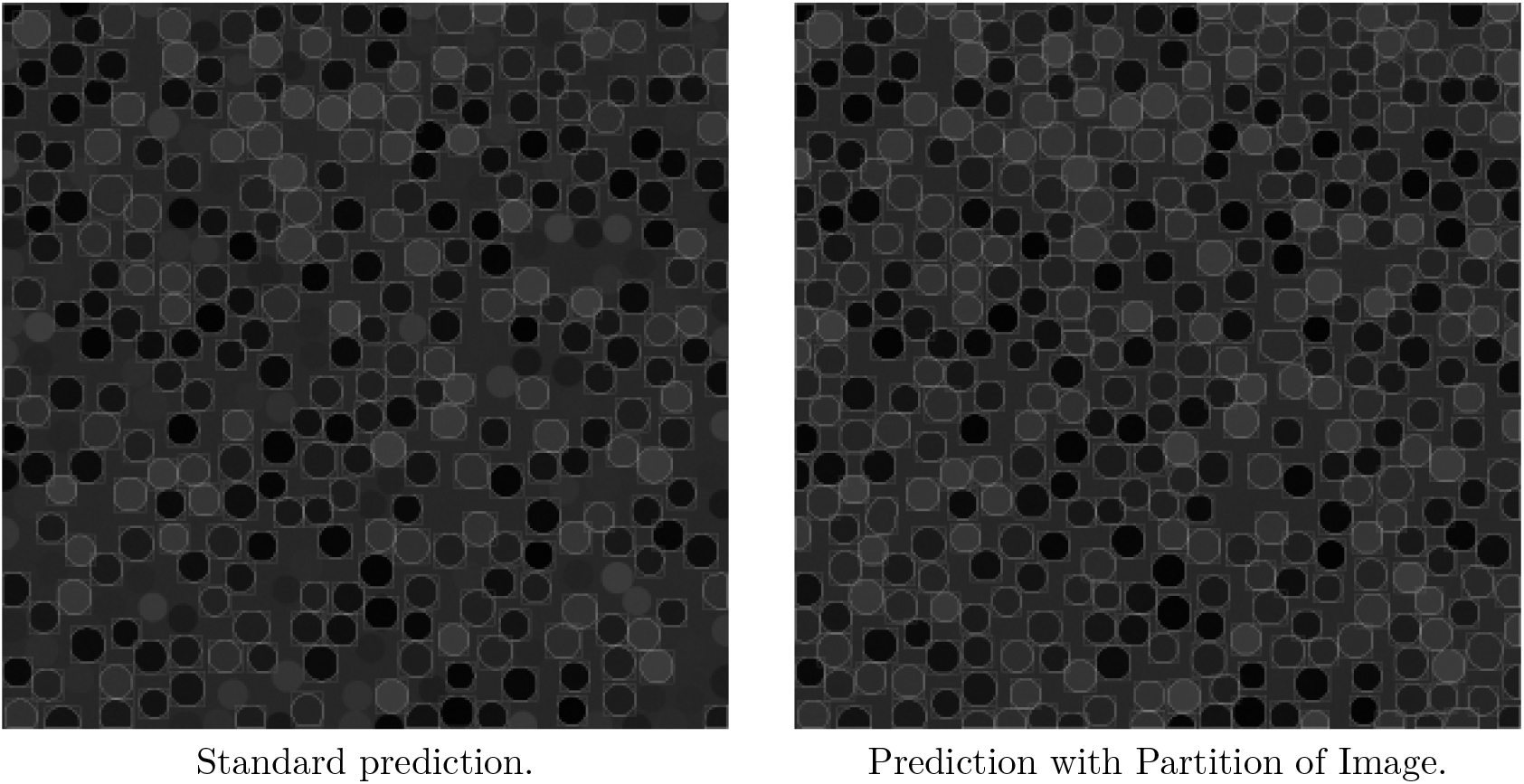
Mask-RCNN predictions (contours and bounding boxes) without and with Partition of Image on an image of original size 2000^2^. The first model finds 286 cells, and the second 397 - the correct number is 400.

**Table 1:**
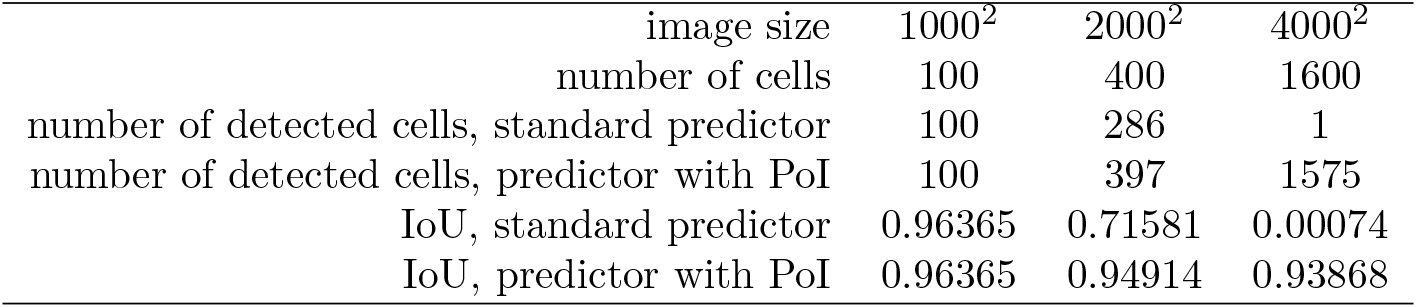
Comparison of detection results from Mask-RCNN model without and with Partition of Image (PoI), according to cell count and Intersection over Union (IoU) metrics. Due to the standard predictor having to resize images of size 2000^2^ and 4000^2^, accuracy on those deteriorate considerably.

### 2.2 Tablet Annotations

The rise of supervised machine learning models in image analysis implies many tasks now need images to be annotated. While annotation is certainly possible using a typical mouse or trackpad, the activity quickly becomes tiresome as that is not the optimal input for free-hand drawing. Fortunately, the popularity of tablets with digital pencil tools (such as the iPad and the Apple Pencil) provide a means to improve the speed and accuracy of annotations.

Since `publicityWebApp.py` uses a web-browser as the user-interface, and can be deployed to a local WIFI network, its annotation tools can be therefore used from a connected tablet. Figure 2 shows a screenshot of the PuBliCity Web-App interface running on an iPad, accessing the web-server running on a PC in the same WIFI network.

**Figure 2:**
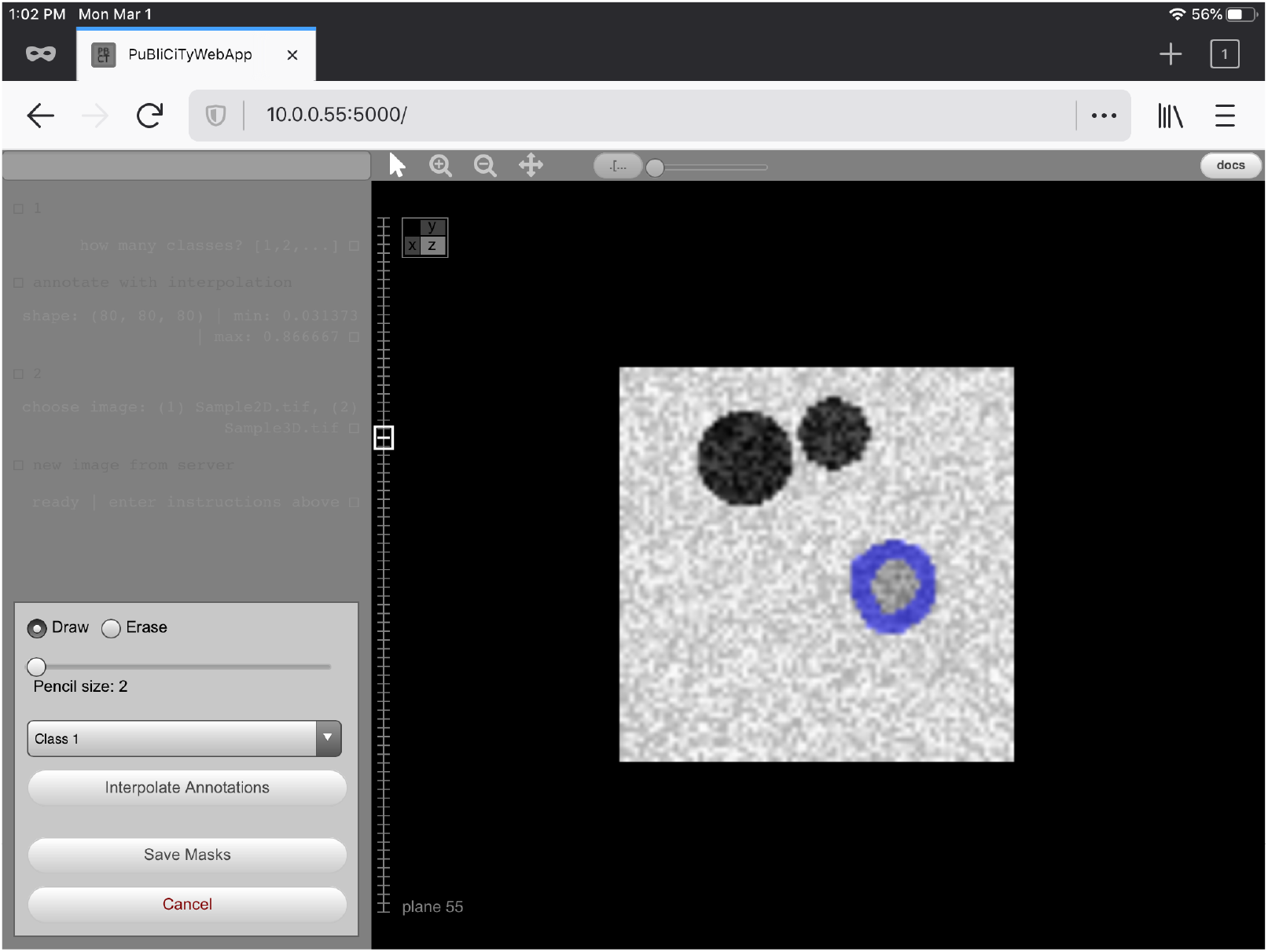
iPad screenshot of the PuBliCiTy Web-App. The user is annotating (blue highlight) a synthetic 3D image using an Apple Pencil.

Besides improving user experience, the web-app’s “annotate with interpolation” functionality is particularly useful in generating 3D annotations, as it interpolates sparsely annotated planes to generate solid 3D blocks of labeled voxels.

## 3 Access and Documentation

We recommend the reader to refer to PuBliCiTy’s GitHub page for up-to-date source code, tutorials, and documentation: github.com/hms-idac/PuBliCiTy.

Most of the functions are documented, and there are currently (i.e. as of this writing) six tutorials, covering the most advanced functionalities, such as pixel and voxel classification (using Random Forests and Deep Learning methods), instance segmentation (via Mask-RCNN), and the PuBliCiTy Web-App.

